# Genome sequencing and multi-stage, blood-feeding, and tissue-specific transcriptome atlas of the Rocky Mountain wood tick provide a critical resource for this vector

**DOI:** 10.64898/2026.04.15.717773

**Authors:** Joshua E. Tompkin, Perot Saelao, Justyna Kruczalak, Huiqing Yeo, Pia U. Olafson, Sheina B. Sim, Kennan Oyen, Melissa Kelley, Renee L. Corpuz, Brian Scheffler, Scott M. Geib, Anna K. Childers, Xiaoting Chen, Matthew T. Weirauch, Shaun J. Dergousoff, John Soghigian, Susan M. Noh, Joshua B. Benoit

## Abstract

*Dermacentor anderson*i, the Rocky Mountain wood tick, is an important vector for pathogens impacting human and animal health, including bovine anaplasmosis, Colorado tick fever, and Rocky Mountain spotted fever. A better understanding of the biology of this tick is needed for developing disease prevention and vector control strategies. A reference genome was assembled for *D. andersoni* using high-fidelity (HiFi) long-read PacBio sequences and Hi-C contact mapping, yielding a contiguous assembly in which most contigs matched one of 11 chromosomes. Genome annotation by the NCBI eukaryotic genome annotation pipeline revealed high gene content completeness, yielding a genome completeness score of 94.0% using the Arachnida ortholog dataset. Following genome sequencing, we identified specific genes involved in blood feeding across a range of tissue types and life stages for *D. andersoni*. To accomplish this, RNA-seq analysis was used to investigate differential gene expression across most organs in adult, nymphal, and larval *D. andersoni* before and after feeding. Based on this analysis, we identified several gene groups that are involved in blood feeding. Furthermore, we establish sex- and developmental-stage-specific transcriptional profiles. Collectively, this study advances knowledge of *D. andersoni* biology and enables the development of strategies to limit the spread of diseases transmitted by this tick.

## Introduction

*Dermacentor andersoni* Stiles, 1908, commonly known as the Rocky Mountain wood tick, is a member of the family Ixodidae and is found primarily in the western United States and southwestern Canada along the Rocky Mountains, and in mixed grassland prairie regions (Dergousoff et al., 2025; Lindquist et al., 2016; Rochon et al., 2012). Like other ixodid ticks, *D. andersoni* has a three-host life cycle, larva, nymph, and adult, with each stage feeding once on a vertebrate host before molting or reproducing. This tick feeds on a broad range of mammals, including humans, livestock, and wildlife, and its prolonged blood-feeding behavior allows the uptake and transmission of pathogens over several days. The tick’s ecology and host-seeking behavior make it not only a key species in North American tick biodiversity but also an important subject for studies of vector biology and disease ecology.

The public health and veterinary significance of *D. andersoni* stems from its role as a vector for multiple pathogens. It is a vector of Rocky Mountain spotted fever (*Rickettsia rickettsii*), Colorado tick fever virus, tularemia (*Francisella tularensis*), and bovine anaplasmosis (*Anaplasma marginale*), and has been implicated in the transmission of Q fever (*Coxiella burnettii*) (Clark et al., 2025; Dergousoff and Chilton, 2012). The ability of this tick to transmit these agents means that human and animal exposures in endemic areas can result in serious, sometimes fatal, illnesses. In addition to pathogenic microbes, *D. andersoni* hosts diverse bacterial endosymbionts that influence tick physiology and susceptibility to infection (Clayton et al., 2015; Niebylski et al., 1997), highlighting complex interactions among the microbiota, tick biology, and disease transmission. Tick paralysis caused by *D. andersoni* is a significant health concern for both humans and animals (Dworkin et al., 1999; Lysyk et al., 2009), resulting from a neurotoxin in the tick’s saliva that can induce progressive paralysis and, in severe cases, respiratory failure if the tick is not promptly removed.

Genomic research on ticks has expanded rapidly over the past decade (Cassens et al., 2025; De et al., 2023; Jia et al., 2020), providing critical insights into the biology, evolution, and vector capacity of these medically important arthropods. Tick genomes are generally large and complex compared to those of many other arthropods, often exceeding 1–2 Gb (Geraci et al., 2007), and are characterized by high levels of repetitive elements, gene duplications, and expanded gene families associated with blood feeding, host sensing, and immune modulation (Cassens et al., 2025; De et al., 2023; Jia et al., 2020). Integration of genome assemblies with transcriptomic, proteomic, and epigenomic data has further enabled stage-, tissue-, and sex-specific analyses of gene expression, illuminating developmental regulation and vector competence. Tick genome and transcriptomic resources provide a robust framework for comparative biological studies on ticks, advancing our understanding of tick–pathogen–host interactions and supporting the development of novel strategies for tick control and prevention of tick-borne diseases.

Here, we report a high-quality genome assembly of *D. andersoni* generated using PacBio HiFi sequence assembly scaffolded into chromosomes using Hi-C contact information and annotated with the National Center for Biotechnology Information (NCBI) eukaryotic genome annotation pipeline, yielding a reference with high contiguity and completeness. Along with the genome, we provide extensive transcriptomic analyses of *D. andersoni* across development, tissues, and blood feeding, providing a comprehensive view of how gene expression is dynamically regulated to support tick growth, survival, and blood ingestion. These transcriptomic analyses, which include RNA-seq experiments across developmental stages and tissues, reveal gene expression patterns associated with feeding, development, and sex-specific biology in *D. andersoni*. These genomic and transcriptomic resources enable researchers to identify genes that define unique traits of this tick, such as blood-feeding mechanisms, chemosensory systems for host detection, and molecular interactions with pathogens.

The availability of the *D. andersoni* genome enhances species-specific research and comparative genomics and transcriptomics across tick vectors, identifying shared and divergent genomic features that underlie pathogen transmission and vector competence. With robust genomic and transcriptomic resources, investigators can explore gene families that mediate host–parasite interactions, characterize salivary proteins that suppress host immune responses, and assess genetic variation across wild populations for *D. andersoni*. Ultimately, the insights from these resources for *D. andersoni* support the development of targeted interventions—such as novel acaricides or vaccine control strategies—to mitigate the spread of tick-borne diseases affecting humans and animals.

## Materials and Methods

### Sample collection

Rocky Mountain wood ticks used for DNA or RNA sequencing originated from laboratory colonies established with specimens collected from the Reynolds Creek watershed in Owyhee County, southwestern Idaho or Dubois, Idaho, respectively. The colonies were maintained following the methods previously described (Scoles et al., 2005; Stiller et al., 2002). The Reynolds Creek colony ticks have been reared in the laboratory for >20 years (about 40 generations). The Dubois colony was reared in the laboratory for 2 years.

### DNA extraction and assembly

Tick samples were flash frozen in liquid nitrogen and shipped to the USDA-ARS Tropical Pest Genetics and Molecular Biology Research Unit in Hilo, Hawaìi, USA, on dry ice for library preparation. This began with high molecular weight genomic DNA from a single *D. andersoni* female using the MagAttract HMW DNA Kit (Qiagen, Hilden, Germany) with the fresh or frozen tissue protocol, without modifications, followed by a 2.0x SPRI bead clean-up to improve sample purity. DNA quantity was determined using the fluorometer feature of a DS-11 Spectrophotometer and Fluorometer (DeNovix Inc, Wilmington, DE, USA) and dsDNA Broad Range (BR) Qubit assay (ThermoFisher Scientific, Waltham, MA, USA). DNA purity was determined using the UV-Vis spectrometer feature on the DS-11. To ensure the genomic DNA was the appropriate size for library preparation, the DNA was sheared to a narrow size distribution with a mean fragment length of approximately 20 kb using a Diagenode Megaruptor 2 according to the manufacturer’s protocol (Denville, New Jersey, USA). Following this, the size distribution of the sheared DNA was determined on a Fragment Analyzer (Agilent Technologies, Santa Clara, California, USA) using the High Sensitivity (HS) Large Fragment kit. The sheared genomic DNA served as the starting input for PacBio SMRTBell library preparation.

The SMRTbell library was prepared using the SMRTbell Express Template Prep Kit 2.0 according to the manufacturer’s protocol (Pacific Biosciences, Menlo Park, California, USA). The final library was shipped to the USDA-ARS Genomics and Bioinformatics Research Unit in Stoneville, Mississippi, USA, where it was bound to the sequencing polymerase and sequenced on one Pacific Biosciences 8M SMRT Cell on a Sequel IIe system (Pacific Biosciences, Menlo Park, California, USA) with a pre-extension time of 2 hours and a collection time of 30 hours. DNA sequencing results are available under the NCBI Project PRJNA767396 (SRR16118521).

In parallel to HiFi library preparation, a Hi-C library was prepared using another *D. andersoni* female from the same colony. Arthropod tissue was crosslinked using the Arima Hi-C low-input protocol, and proximity ligation was performed using the Arima Hi-C Kit (Arima Genomics, San Diego, California, USA). Afterwards, the DNA was sheared using a Diagenode Bioruptor Pico and then size-selected to enrich for DNA fragments from 200-600bp. An Illumina library was prepared from the sheared and size-selected DNA using the Swift Accel NGS 2S Plus kit (Integrated DNA Technologies, Coralville, Iowa, USA) and sequenced on a NovaSeq 6000 at the Hudson Alpha Genome Sequencing Center (Huntsville, Alabama, USA). Hi-C results are available under the NCBI Project PRJNA767396 (SRR18268591).

Adapter-contaminated reads in all CCS (HiFi) datasets were identified using NCBI FCS-adaptor (Astashyn et al., 2024) and then removed from the read pool using HiFiAdapterFilt (Sim et al., 2022). Following this, the HiFi reads were assembled into contigs using HiFiASM v0.19.3-r572 (Cheng et al., 2021). The primary and alternate libraries served as the input to the PurgeDups pipeline (Guan et al., 2020) to remove duplicate regions or contigs. Paired Hi-C reads were mapped to the duplicate purged primary assembly using BWA-mem2 (Vasimuddin et al., 2019), and the mapped reads were used to generate a Hi-C contact map following the Hi-C YAHS pipeline (Zhou et al., 2023). Hi-C contacts were edited to ensure correct match using Juicebox v2.11 (Durand et al., 2016). Genome annotation was performed using the NCBI Eukaryotic Genome Annotation Pipeline (EGAP) (Thibaud-Nissen et al. 2013). The genome and Refseq data sets are available through the NCBI (GCA_023375885.3 and GCF_023375885.2)

### Genome quality assessment

The *D. andersoni* genome assembly was assessed for completeness using the Arachnida v10 database (odb10) of the BUSCO v5.2.2 software (Tegenfeldt et al., 2025). Taxonomic assignment of duplicate purged contigs was determined by simultaneously aligning contigs to the NCBI non-redundant nucleotide database (released 2022-06-01) using the blastn function of Basic Local Alignment Search Tool (BLAST) + (Camacho et al., 2009) and aligning contigs to the UniProt protein database (released 2020-06-17) using the blastx function of Diamond (Buchfink et al., 2021). Local alignment results were summarised using the Blobtools2 v4.1.5 pipeline (Challis et al., 2020) and Blobblurb v2.0 (Sim, 2022). Off-target contigs and sequences were also identified using NCBI FCS-GX, a cross-species aligner that classifies sequences as contaminants when their taxonomic assignment is different from the provided taxonomic identifier.

Mitochondrial contigs were identified using MitoHiFi v3.0.0 (Uliano-Silva et al., 2023) using the complete mitochondrion genome of *D. andersoni* (NCBI RefSeq accession: NC_061057). Following taxonomic assignment of contigs, mitochondrial contig detection, and Hi-C scaffolding, non-arthropod and duplicate mitochondrial contigs were removed from the remaining contigs that could not be placed into chromosomes based on Hi-C contact data.

Putative tRNA genes were identified by genome analysis using tRNAscan-SE 2.0 with parameters optimized for arthropods (Chan et al., 2021; Kelley et al., 2025; Ryazansky et al., 2024). To ensure high-confidence predictions, a minimum quality score threshold of 50 (default = 20) was applied, and searches were restricted to eukaryotic tRNAs with output configured to report predicted tRNA secondary structural features. Predicted tRNAs from this initial screen were subsequently filtered using the recommended Eukaryotic Confidence Filter under default settings to remove tRNA-derived repetitive elements and other non-functional predictions. The resulting dataset represents a high-confidence set of tRNA genes associated with specific amino acids for each species and was used for all downstream comparative analyses.

### Lifestages and tissues for RNA sequencing

All developmental stages and organs from fed and unfed ticks were collected for RNA sequencing. Nymphs were fed to repletion on rats, while all other life stages fed on calves (University of Idaho IACUC-2022-26). Adult males were fed 7 days and adult females fed 5 days prior to forced removal. After feeding, adult male and female ticks were held for 5 days in 98% humidity at 26° C and then dissected. Midguts, salivary glands, the synganglion and Haller’s organ were dissected from unfed and fed adult ticks. Reproductive organs were dissected only from fed adult ticks. Pools of approximately 50 larvae and 50 nymphs in replicates of three were stored. For adult ticks, five organs per tube, in triplicate, were stored. The exception was Haller’s organ, for which 20 organs from male ticks and 20 organs from female ticks were pooled for downstream analysis. Late embryonated eggs were stored in RNALater (Sigma-Aldrich, St. Louis, MO). All other organs and life stages were flash frozen in liquid nitrogen.

### RNA extraction, processing, and sequencing

Following removal from liquid nitrogen, tissues were pulverized using a mortar and pestle. Total RNA was extracted using the Quick-RNA Microprep Kit (Zymo Research, Irvine, CA, USA) following the manufacturer’s protocols. RNA from midguts was cleaned using the RNA Clean and Concentrator-5 Kit (Zymo Research, Irvine, CA, USA). After total RNA extraction, residual genomic DNA was removed using RNase-free DNase I (Zymo Research, Irvine, CA, USA). RNA quantity and purity were assessed spectrophotometrically, and RNA integrity was evaluated by electrophoresis. Polyadenylated mRNA was then enriched from total RNA using oligo(dT)-based selection. RNA-Seq libraries were constructed using the NEBNext library preparation kit (New England Biolabs, Ipswich, MA, USA) following the manufacturer’s protocols. Briefly, purified mRNA was fragmented and reverse-transcribed to generate first- and second-strand cDNA, which was subsequently end-repaired, A-tailed, and ligated to indexed Illumina sequencing adapters. Libraries were amplified by limited-cycle PCR and purified to remove adapter dimers and size-selection contaminants. Final library quality and concentration were verified prior to sequencing. Libraries were pooled in equimolar amounts and sequenced on an Illumina NextSeq 2000 platform to generate paired-end reads of 150 nt, yielding approximately 50-60 million reads per library. All RNA-seq sets are available through the NCBI (Bioproject PRJNA936406).

### RNA-seq quality assessment and analysis

RNA-seq data were analysed using methods similar to those employed in other arthropod systems (Attardo et al., 2019; Olafson et al., 2021; Pathak et al., 2022; Sterkel et al., 2025). The quality of RNA-seq reads was examined using FastQC v0.11.9. Based on FastQC reports, adapter trimming, polyG tail removal, and quality filtering were performed using fastp v0.23.3 (Chen et al., 2018). Only reads containing 50 or more bases were kept for downstream analysis. After trimming, FastQC was used again to ensure that reads were of sufficient quality. HISAT2 was used to map the reads onto the reference genome from NCBI RefSeq GCF_023375885.1 (Kim et al., 2019). The featureCounts program from the Subread package v2.0.3 was used to determine the number of reads that mapped to each exon feature within the genome (Liao et al., 2014). A count matrix containing the number of times each gene was mapped within each sample was created, and statistical analysis of differentially expressed genes was done using the DESeq2 package v1.48.0 in R v4.3.0 (Love et al., 2014). Default settings were used, other than a significance cutoff of 0.05 was used to determine genes that were significantly differentially expressed. Comparisons were done among samples that differed only in tissue, development stage, feeding status, or sex. Genes with a log base two-fold change of two or greater (increase or decrease) were used for further gene ontology analysis. Principal component analysis was used to reduce dimensionality and visualize clustering patterns in gene expression across samples. To obtain a list of significant Gene Ontology (GO) terms associated with the differentially expressed genes for each comparison, the g:GOSt function in g:Profiler was used with default settings (Kolberg et al., 2023). To reduce list size and redundancy of GO terms, clustering and visualization were performed with REVIGO (Supek et al., 2011).

A comparative analysis of the genomes of *D. andersoni, D. silvarum, Galendromus occidentalis, Ixodes scapularis, Ornithodoros turicata, Panonychus citri, Rhipicephalus microplus. R. sanguineus, Tetranychus utricae,* and *Varroa destructor* was performed to determine the presence and expression patterns of genes undergoing rapid expansion in *D. andersoni*. First, redundant sequences arising from alternative splicing were removed across all species by isolating the longest isoform sequence for each gene. Then, OrthoFinder v2.4.4 was used to identify gene families within the species and to construct a rooted species tree for all species analyzed (Emms and Kelly, 2019). CAFE5 v1.1 was used to estimate the rate of expansion or contraction of gene families in *D. andersoni (Mendes et al., 2021)*. The expression patterns of the genes within the significantly expanded gene families of *D. andersoni* were then analyzed using the normalized gene counts from DESeq2.

Weighted gene co-expression network analysis (WGCNA) was performed in R to identify groups of genes with coordinated expression patterns across the sampled conditions and to summarize these patterns at the module level (Langfelder and Horvath, 2008). Normalized expression values generated during the DESeq2 workflow were used as the input matrix for network construction. Before analysis, genes exhibiting no variation across samples were removed to avoid spurious correlations and to reduce noise in downstream module detection. Following this filtering step, 31,290 genes remained and were carried forward for network inference. A signed co-expression network was then constructed by calculating pairwise gene–gene correlations across all samples, transforming the correlation matrix into an adjacency matrix using soft thresholding, and subsequently deriving a topological overlap matrix (TOM) to capture shared neighborhood structure among genes. The selection of the soft-thresholding power (β) was guided by examination of the scale-free topology fit index across a range of candidate powers. Because a scale-free topology is a common empirical property of biological networks, the β value was chosen to balance improved scale-free fit with preservation of meaningful connectivity. Based on these diagnostics, a soft-thresholding power of 5 was selected for network construction. Modules of co-expressed genes were identified using hierarchical clustering of TOM-based dissimilarity followed by dynamic tree cutting, with the minimum module size set to 20 genes to ensure that detected modules represented stable, interpretable clusters rather than very small, potentially noisy groupings.

### Identification of transcription factors and their binding sites

Putative transcription factors (TFs) were identified by screening the predicted protein-coding gene set for conserved DNA-binding domains (DBDs), using a previously established method that has successfully identified TF genes in other arthropod systems (Benoit et al., 2016; Scott et al., 2020; Shippy et al., 2024). Amino acid sequences were scanned using the 81 Pfam models corresponding to known eukaryotic DBDs as previously described (Weirauch and Hughes, 2011) and implemented with the HMMER tool, applying recommended significance thresholds (per-sequence E-value < 0.01; per-domain conditional E-value < 0.01). Proteins containing one or more DBDs were classified into TF families based on the type and arrangement of their domains within the sequence (e.g., bZIP×1, AP2×2, Homeodomain+POU).

For each TF family, DBD sequences were aligned using Clustal Omega with default parameters. In proteins containing multiple DBDs, each domain was aligned independently. Pairwise sequence identity was then calculated for all aligned DBDs, defined as the percentage of identical amino acid residues across the alignment. Predicted DNA-binding specificities were assigned by transferring known specificities from experimentally characterized TFs when sequence identity exceeded previously established family-specific thresholds. For example, the DBD of g19927.t1 shares 87.7% sequence identity with the *Drosophila melanogaster* Antennapedia (Antp) protein; because the homeodomain family threshold is 70% and Antp binding specificity has been experimentally determined, g19927.t1 was inferred to possess the same DNA-binding specificity. Enriched transcription factor binding motifs were detected within the 500-bp and 2,000-bp upstream regions of predicted transcription start sites using the HOMER motif discovery tool, following approaches previously applied to investigate relationships between transcription factors and gene expression dynamics in insects (Rotenberg et al., 2020; Ryazansky et al., 2024; Scott et al., 2020; Shippy et al., 2024).

### Sex chromosome identification

For identification of the sex chromosome in *D. andersoni*, adults from two colonies, as well as their nymph offspring, were chosen from a separate sequencing experiment. The two laboratory colonies are maintained at the Lethbridge Research and Development Centre, Agriculture and Agri-Food Canada. The first colony was established from collections made near Waterton Lakes National Park, Alberta, in 2022. The second colony was established from collections made in 2019 near Quilchena in southern British Columbia. F1 individuals from the first colony were crossed with F3 individuals from the second in 2023 for other projects, resulting in nymph offspring. The tick feeding for the rearing and mating experiment was approved by the Lethbridge Research and Development Centre Animal Care Committee; Animal Use Protocols 2215 and 2217. Parents and nymphs were stored at -20°C and then quadrisected with a sterile scalpel prior to DNA extraction with the E.Z.N.A Forensic DNA kit (Omega Bio-Tek, Norcross, GA, USA). Two library preparations were done for each tick using a multiplex library prep kit (Twist BioScience, San Francisco, CA, USA) at the Centre for Health Genomics and Informatics (CHGI), University of Calgary. The libraries from each sample were sequenced separately on two lanes of the Illumina NextSeq 2000 P4 flow cell.

Resulting sequencing reads were trimmed with fastp version 0.23.4 (Chen et al., 2018) and aligned to a *D. andersoni* reference genome (GCA_023375885.3) with bowtie 2 version 2.4.5 (Langmead and Salzberg, 2012). The bam files for each sample were merged using Samtools version 1.15 (Danecek et al., 2021). Coverage per chromosome was obtained with QualiMap version 2.2.1 (Okonechnikov et al., 2016). Seven adult males, seven adult females, and six nymph offspring were selected for additional analysis to determine the sex chromosome (see supplemental data files for colony identity, sex, and coverage values). Due to differences in coverage among samples, the relative coverage for each chromosome was calculated by dividing the coverage for each chromosome by the average coverage of the other ten chromosomes, excluding CM094977. As the sex of an adult corresponded to differences in relative coverage on CM094977, and this coverage difference was reflected in nymphs, nymphs were also coded as male or female. An ANOVA was used to evaluate whether there were significant differences in coverage of the putative sex chromosome across life stages and sexes. Then, relative coverage across chromosomes was visualized via a violin plot (Fig. 11).

## Results and Discussion

### Genomic and transcriptome summaries

Assembly completeness was assessed using Benchmarking Universal Single-Copy Orthologs (BUSCO) with the arthropoda_odb9 dataset. BUSCO analysis identified 94% of expected arthropod orthologs as complete in the assembly, indicating that the majority of conserved gene content is represented (Fig. 1). This level of completeness is consistent with high-quality arthropod genome assemblies, specifically other tick species (Cassens et al., 2025; De et al., 2023; Jia et al., 2020), and suggests minimal loss of genic regions during assembly. The Hi-C contact map showed strong diagonal interaction signals, consistent with well-resolved chromosome-scale scaffolds, resulting in a contiguous assembly in which most contigs matched one of the 11 chromosomes. Limited off-diagonal interaction signals were observed, indicating few misjoins and supporting the accuracy of scaffold placement. The combination of high contiguity metrics, substantial BUSCO completeness, and well-defined Hi-C interaction patterns demonstrates that this genome assembly is of high quality. Importantly, these metrics are comparable to those reported for other recently sequenced tick species with chromosome-level assemblies (Cassens et al., 2025; De et al., 2023; Jia et al., 2020), supporting the utility of this genome as a robust resource for comparative genomics, evolutionary analyses, and studies of tick biology and vector competence. tRNA gene expansion was noted from *D. andersoni* and other ticks (Fig. 2, Table S1), with over 20-30 fold increases in tRNA number compared to other members of Acari and insects. A higher number of tRNA genes did not occur in tRNAs for specific amino acids and isotypes among ticks. Rather, there was a uniform increase across the entire tRNA repertoire. Notably, the magnitude of this expansion is substantially greater than that observed in Culicinae mosquitoes relative to other dipteran flies (Ryazansky et al., 2024), highlighting a lineage-specific amplification of translational capacity in ticks, which may be linked to their large genome sizes (Geraci et al., 2007; Jia et al., 2020)

**Figure 1.**
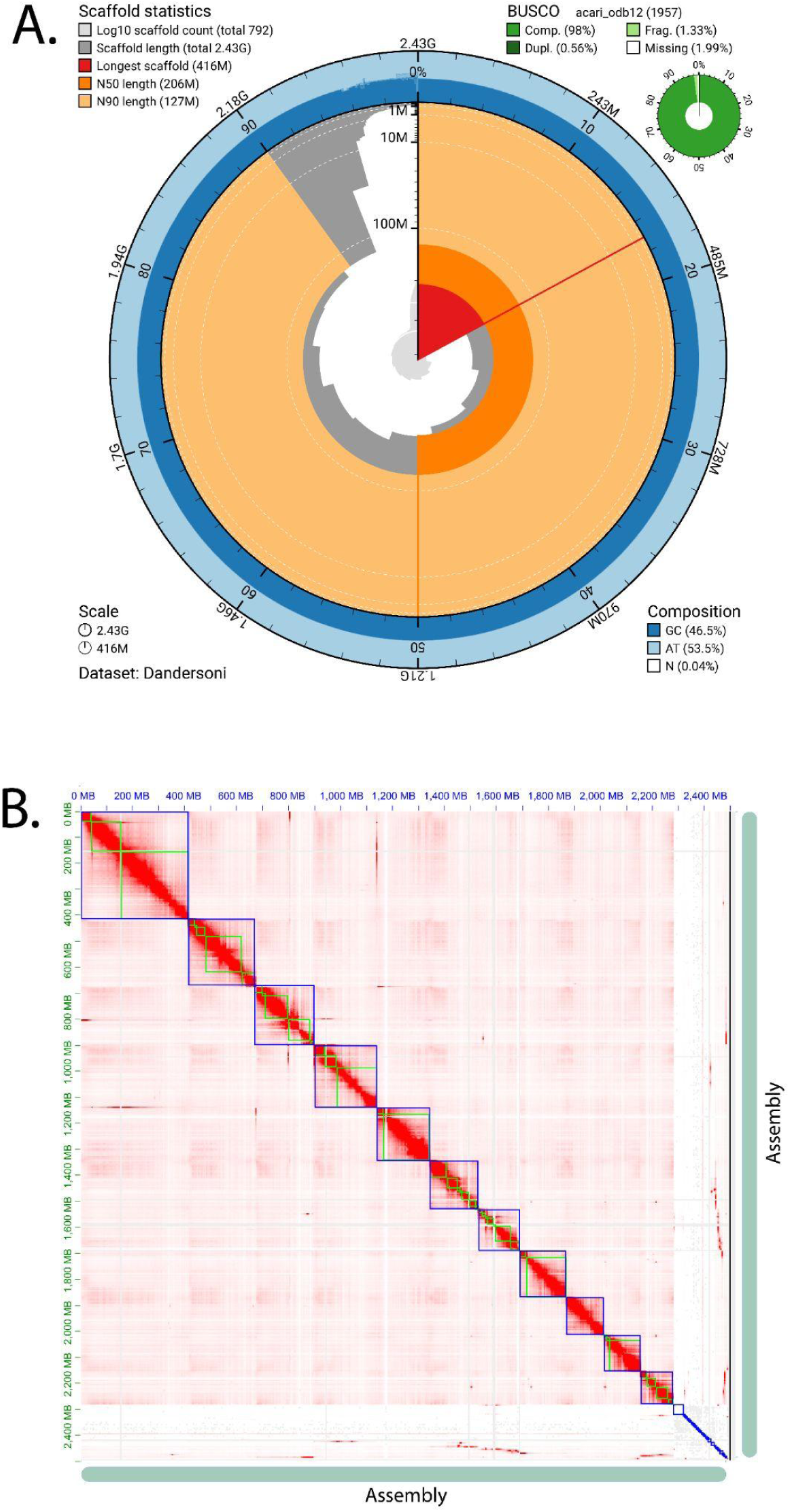
Summary of *D. andersoni* genome assembly. (A) Snail plot showing contig length statistics and BUSCO analysis results. The axis around the circle indicates the proportion of the total assembly composed of contigs at least as long as the length shown on the vertical axis. Statistics, including the longest contig, N50, and N90, are highlighted in color. The top-right plot shows the results of the BUSCO analysis: the percentages of complete, single-copy, complete-and-duplicate, fragmented, and missing orthologs. (B) Hi-C chromatin contact map showing the frequency of chromatin interactions throughout the *D. andersoni* genome assembly. The position within the assembly to which the first and second reads of each read pair map is shown on the x- and y-axes, respectively. The color of each square indicates the number of read pairs mapped to that location in the assembly.

**Figure 2.**
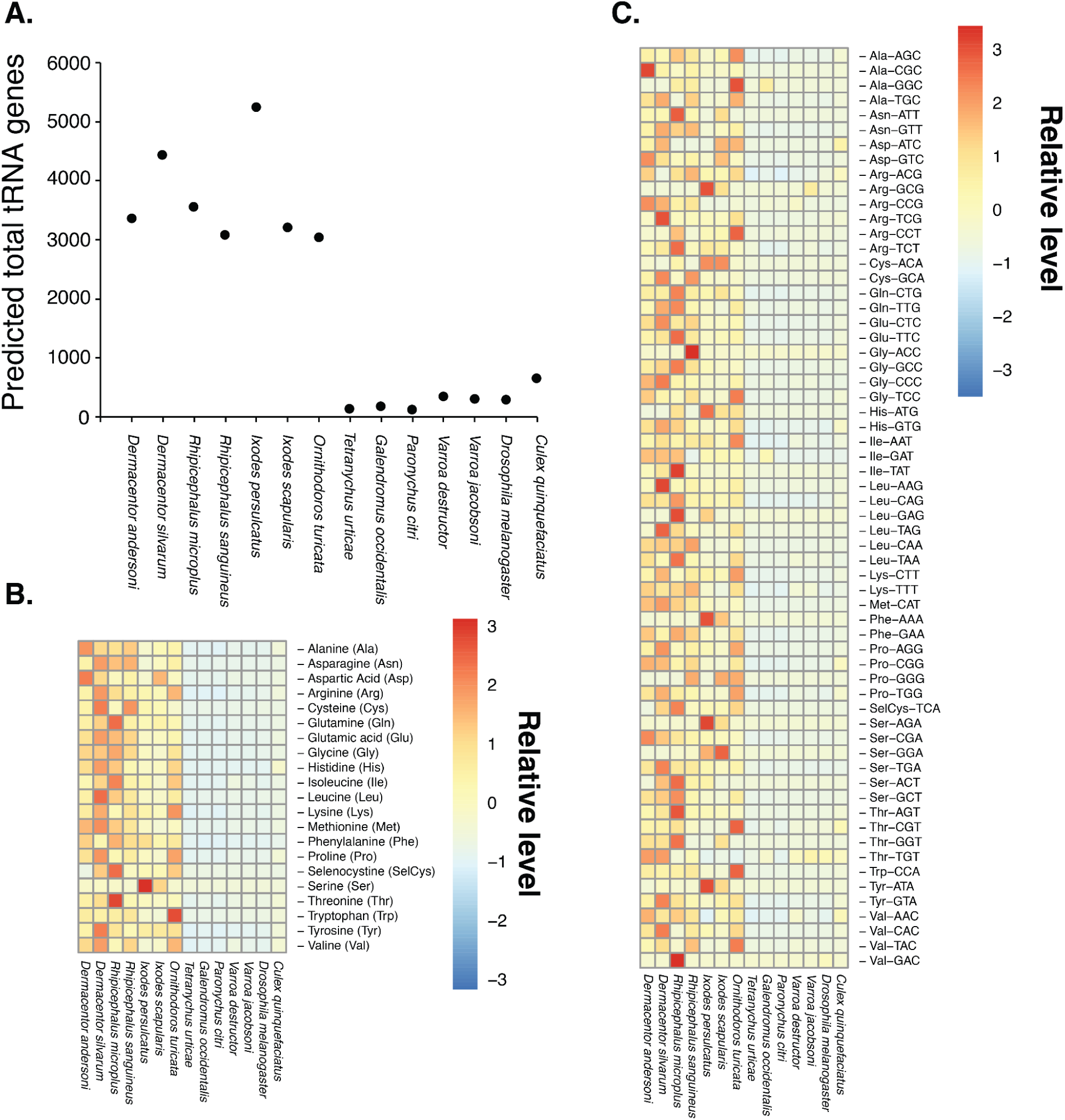
tRNA content of *D. andersoni* compared to other ticks and mites. (A) Changes in the number of predicted tRNA genes associated with individual amino acids. (B) Comparison of predicted tRNA gene repertoires in *D. andersoni* relative to other ixodid ticks, argasid ticks, and representative mite species. (C) Differences in tRNA isotype distributions in *D. andersoni* compared with other ticks and mites. (D) Differences in tRNA anticodon usage in *D. andersoni* relative to other ticks and mites. Specific results are in Table S1.

Extensive RNA-seq studies were used to assess developmental stage, tissue type, sex, and blood-feeding status. All samples were of high quality. Principal component analysis (PCA) based on DESeq2 analysis revealed clear, biologically meaningful structuring of tick samples by developmental stage, tissue type, sex, and blood-feeding status (Fig. 3). Samples clustered strongly by developmental stage along the primary principal components, indicating that ontogenetic transitions drive global transcriptional variation in ticks. Within developmental stages, additional separation was observed among tissue types, consistent with distinct tissue-specific transcriptional programs associated with functions such as digestion, reproduction, and host interaction (Kotsyfakis et al., 2015; Medina et al., 2022; Tirloni et al., 2020). Sex-specific differences were evident, with male and female samples forming discrete clusters, reflecting divergent reproductive physiology and sex-biased gene expression (Edwards et al., 2025; Meibers et al., 2019; Tidwell et al., 2024). Notably, blood feeding produced pronounced shifts in PCA space, with fed and unfed ticks separating along a central axis of variation, highlighting the extensive transcriptional remodeling that accompanies hematophagy in ticks (Kotsyfakis et al., 2015; Medina et al., 2022; Reyes et al., 2024). More details are provided below based on specific biological comparisons related to the blood feeding, tissue, and developmental dynamics. Along with the DESeq2-based analysis, we have conducted WGCNA across developmental stages and tissues, providing a secondary analysis of genes with specific expression profiles beyond the developmental and tissue analyses discussed in the remaining results (Fig. S1, Supplemental Material 1). Together, these patterns demonstrate that tick gene expression is shaped by life history, tissue specialization, sexual dimorphism, and feeding status, and that these factors collectively account for a substantial proportion of the observed transcriptomic variation.

**Figure 3:**
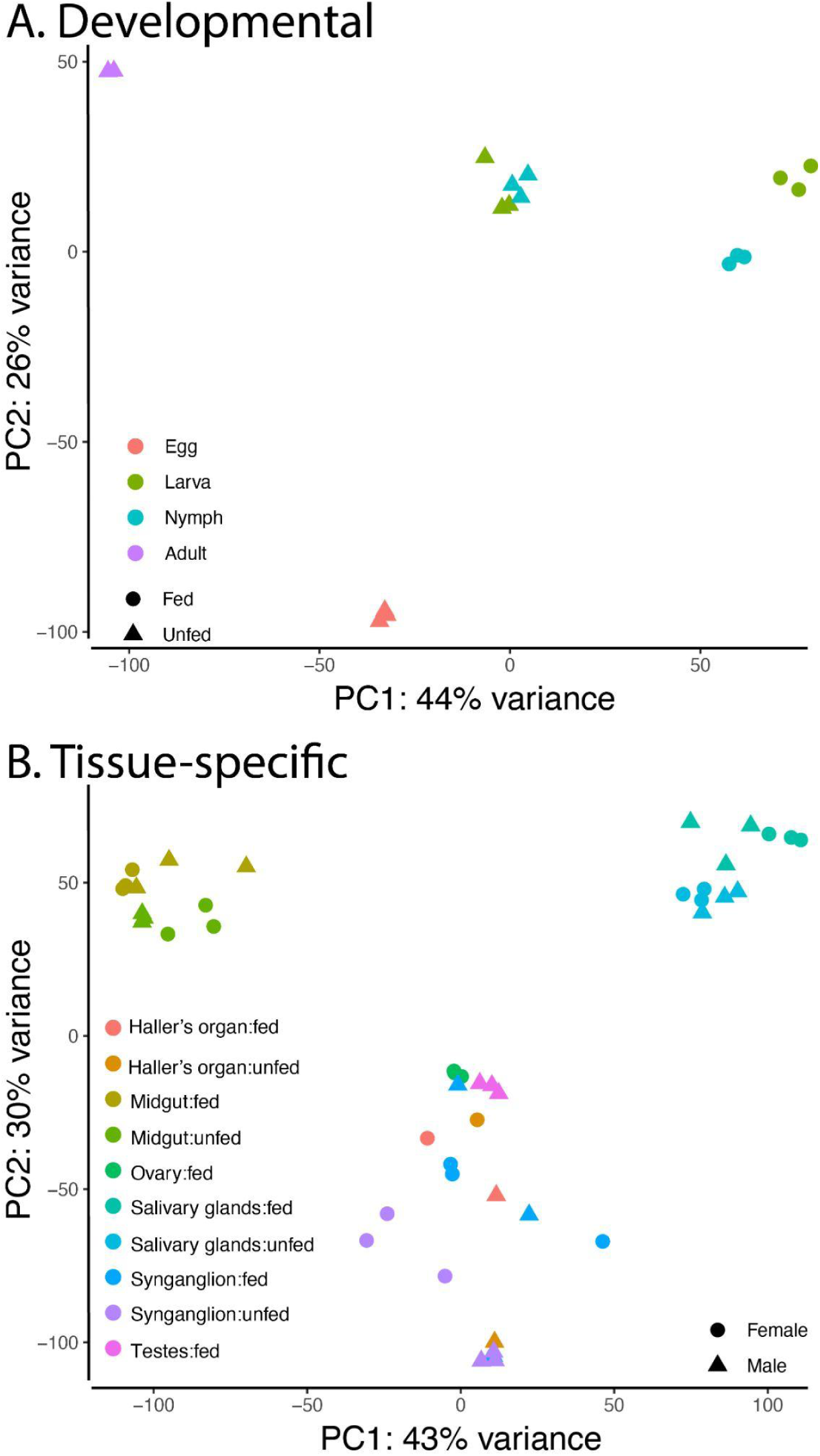
Principal component analysis (PCA) of *D. andersoni* transcriptomes. PCA was performed on normalized RNA-seq expression data based on DESeq2 to visualize global patterns of transcriptional variation across developmental stages, tissues, sex, and feeding status in *D. andersoni*. (A) Developmental PCA showing clustering of samples from egg, larval, nymphal, and adult stages. Symbols denote feeding status (fed vs. unfed), and colors represent developmental stages. Principal component 1 (PC1) explains 44% of the total variance, while PC2 explains 26%, indicating strong stage-specific transcriptional differentiation. (B) Tissue-specific PCA illustrating separation of transcriptomes from Haller’s organ, midgut, ovary, salivary glands, synganglion, and testes under fed and unfed conditions. Symbol shape indicates sex (female or male), and colors denote tissue and feeding state. PC1 and PC2 account for 43% and 30% of the variance, respectively, and demonstrate distinct clustering by tissue type, with additional modulation by sex and blood-feeding status.

### Developmental transcript expressional shifts

To assess transcriptional changes across developmental stages, we conducted RNA-seq analysis of adults, nymphs, larvae, and eggs. Eggs exhibited the largest set of uniquely expressed transcripts compared with later stages, consistent with intense molecular activity associated with embryogenesis (Fig. 4, Table S2). Functional enrichment analyses revealed overrepresentation of DNA-templated transcription, RNA metabolic processes, macromolecule biosynthetic pathways, and developmental signaling pathways, including Wnt and cell-surface receptor signaling. These patterns reflect active cell proliferation, patterning, and differentiation processes required for early development. Similar enrichment of transcriptional and developmental pathways has been reported in eggs of other hard ticks, including *I. ricinus* and *R. turanicus* (Edwards et al., 2025; Ruiling et al., 2022; Vechtova et al., 2020), suggesting a conserved embryonic transcriptional architecture for ticks. Larval and nymphal stages displayed both shared and stage-specific expression profiles, both before and after feeding. Each stage was characterized by enrichment of genes involved in translation, ribosome biogenesis, amino acid activation, and primary metabolic processes, consistent with differentiation following hatching and molting into active stages (Ruiling et al., 2022; Vechtova et al., 2020).

**Figure 4:**
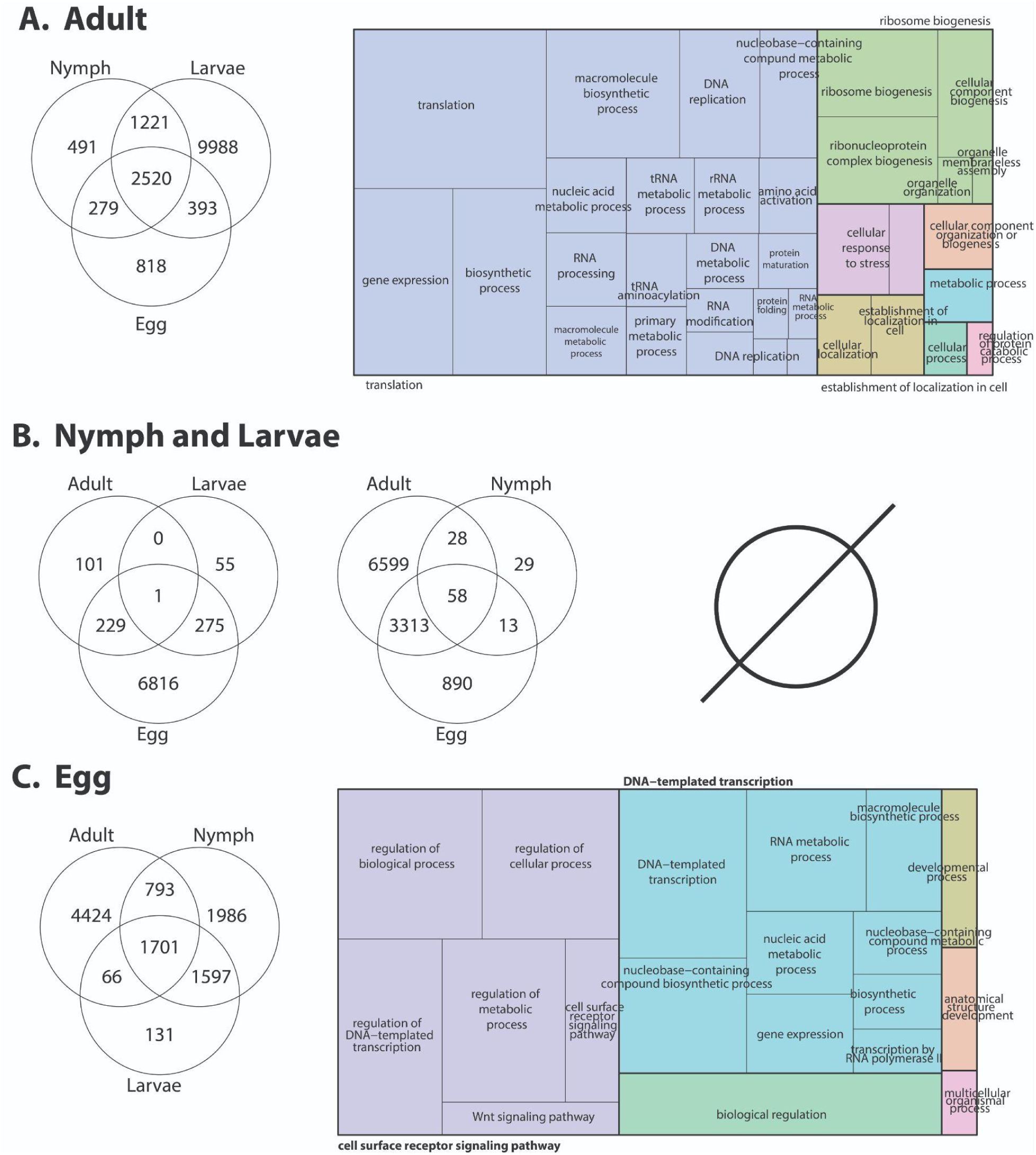
Developmental stage–specific transcriptomic profiles of unfed *D. andersoni*. RNA-seq differential expression analyses were conducted on unfed *D. andersoni* across four developmental stages (A. adult, B. nymph and larva, and C. eggs). Venn diagrams illustrate the number of differentially expressed genes (DEGs) that are unique to or shared among developmental stages, highlighting both stage-specific and conserved transcriptional signatures during tick development in the absence of blood feeding. Nymphs and larvae have no enriched Gene Ontology (GO) terms, indicated by the null symbol. Adults and eggs have enriched GO categories shown as treemaps on the right of the venn diagrams based on the biological processes associated with DEGs in each stage. Complete results for egg and adults are in Table S2 and S3, respectively.

Unfed adults exhibited a distinct expression profile compared with immature stages, with enrichment of genes involved in gene expression regulation, protein folding and maturation, cellular localization, and ribonucleoprotein and organelle biogenesis (Fig. 4, Table S3). These patterns suggest a shift from growth-focused programs toward maintaining complex tissues and physiological readiness for reproduction and host-seeking (Edwards et al., 2025; Ruiling et al., 2022; Vechtova et al., 2020). Notably, despite the absence of blood feeding in these samples, adult ticks already display transcriptional investment in pathways that support cellular homeostasis and responsiveness, consistent with observations in unfed adult ticks, where preparatory transcriptional states precede feeding (Edwards et al., 2025; Rosendale et al., 2019; Ruiling et al., 2022; Vechtova et al., 2020).

Together, these results demonstrate that *D. andersoni* development is accompanied by pronounced and orderly shifts in gene expression programs. The close correspondence between the developmental transcriptional patterns observed here and those reported in other tick species suggests that many of these regulatory programs are evolutionarily conserved (Edwards et al., 2025; Rosendale et al., 2019; Ruiling et al., 2022; Vechtova et al., 2020). At the same time, the magnitude and specificity of stage-enriched expression underscore the importance of considering developmental context when interpreting tick transcriptomes and identifying candidate genes relevant to physiology, vector competence, and control strategies.

### Gene expression changes following blood feeding in larvae and nymphs

In the unfed state, larvae and nymphs were characterized by enrichment of metabolic and biosynthetic processes (Fig. 5, Table S4), including amino acid metabolism, nucleotide and small-molecule metabolism, DNA replication, and protein folding (Guo et al., 2019; Hu et al., 2020; Ruiling et al., 2022). These pathways are consistent with baseline physiological demands associated with pre-feeding questing and survival (Rosendale et al., 2019, 2016; Ruiling et al., 2022). Similar enrichment of core metabolic processes in unfed immature stages has been reported (Guo et al., 2019; Hu et al., 2020; Ruiling et al., 2022), suggesting a conserved metabolic foundation that supports tick development before host attachment.

**Figure 5:**
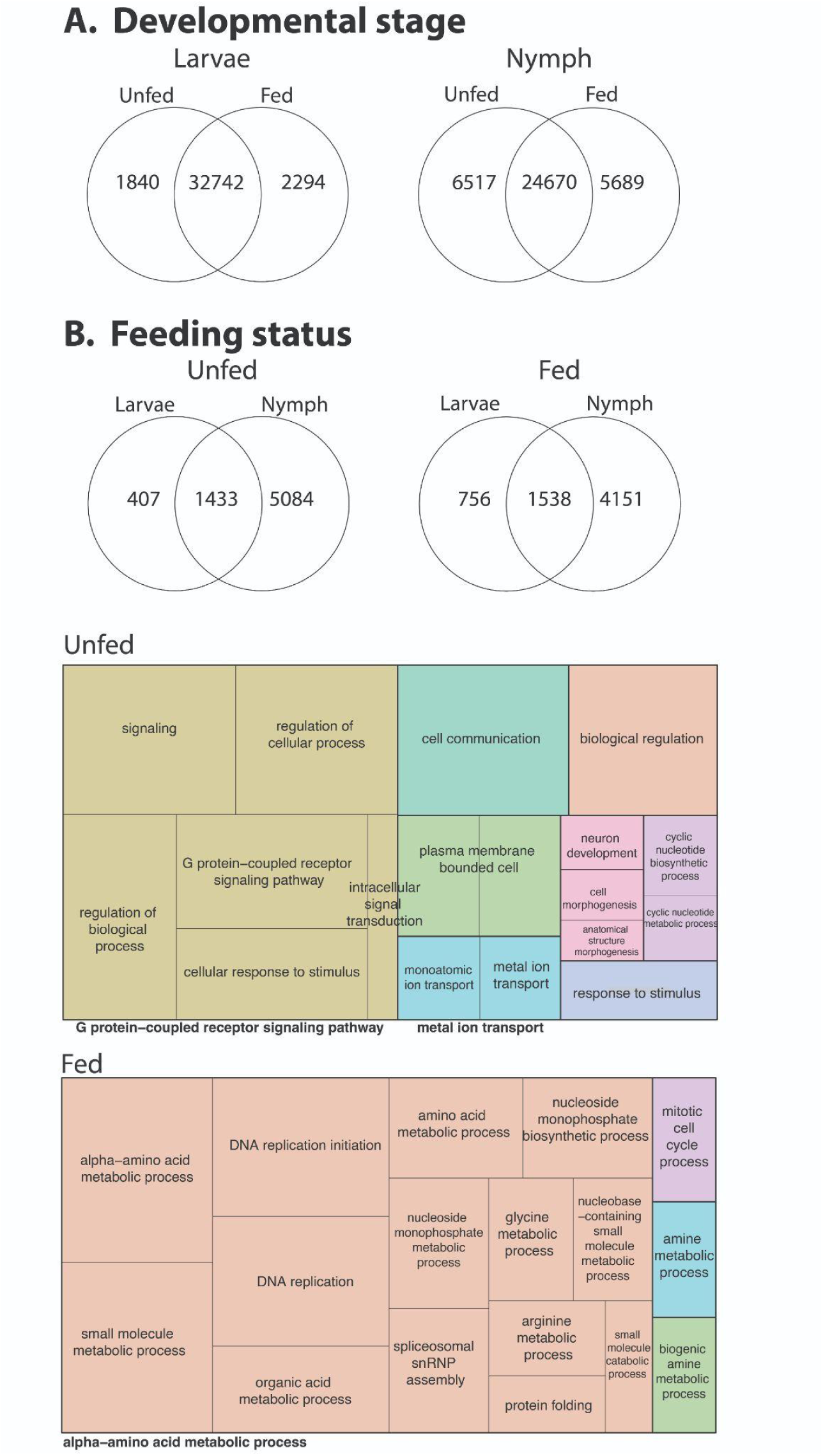
Transcriptomic responses of larval and nymphal *D. andersoni* to blood feeding. RNA-seq differential expression analyses were performed on unfed and fed *D. andersoni* larvae and nymphs. Venn diagrams depict the number of differentially expressed genes (DEGs) that are unique to or shared between larvae and nymphs before and after blood feeding (A and B), illustrating both stage-specific and conserved transcriptional responses to the blood meal. Treemap visualizations summarize enriched Gene Ontology (GO) biological processes associated with DEGs in unfed and fed states. Complete results for fed and unfed ticks are in Table S4 and S5, respectively.

Blood feeding triggered a dramatic shift in gene expression profiles in both larvae and nymphs, with strong enrichment of signaling- and response-related pathways (Fig. 5, Table S5). In fed ticks, differentially expressed genes (DEGs) were associated with G protein–coupled receptor signaling, intracellular signal transduction, cyclic nucleotide biosynthesis and metabolism, ion and metal transport, and cellular responses to stimuli. Additional enrichment of pathways linked to cell communication, morphogenesis, and neuronal development suggests that blood feeding initiates broad physiological coordination across tissues, likely reflecting sensory processing, nutrient sensing, and systemic regulation required during hematophagy. Comparable feeding-induced activation of signaling, ion transport, and neuroregulatory pathways has been documented in fed nymphs and larvae of other tick species (Guo et al., 2019; Hu et al., 2020; Ruiling et al., 2022), underscoring the conserved nature of these responses across hard ticks.

Collectively, these results demonstrate that blood feeding acts as a central transcriptional switch in immature *D. andersoni*, overriding baseline developmental programs and activating signaling- and response-driven gene networks. The strong parallels between *D. andersoni* and other tick species indicate that many blood-feeding–responsive pathways are evolutionarily conserved (Guo et al., 2019; Hu et al., 2020; Ruiling et al., 2022). At the same time, the observed stage-specific differences highlight how developmental context shapes the transcriptional response to blood feeding. These findings emphasize the importance of the larval and nymphal stages as dynamic, biologically active phases of the tick life cycle and provide a framework for identifying feeding-associated genes that may contribute to pathogen acquisition and transmission during these stages.

### Tissue-specific expression changes following blood feeding in males and females

We also observed transcriptional remodeling of the *D. andersoni* midgut in response to blood feeding (Fig. 6, Table S6-7), with patterns that closely mirror findings from midgut RNA-seq studies in other hard tick species (Anderson et al., 2008; Lu et al., 2024; Medina et al., 2022). In both females and males, blood feeding induces shifts in gene expression associated with amino acid, carbohydrate, nucleotide, and small-molecule metabolic processes, reflecting the midgut’s central role in digestion and nutrient assimilation following a blood meal. The enrichment of lipid localization and transport processes aligns with established evidence that the tick midgut serves as a key hub for lipid trafficking, including sterols and cholesterol that must be acquired from the host and redistributed systemically (Anderson et al., 2008; Lu et al., 2024, 2023; Medina et al., 2022). Comparisons with salivary gland and synganglion-associated gene sets further emphasize the tissue specificity of the feeding response, with the midgut dominated by metabolic and biosynthetic functions rather than secretory or neuroregulatory processes, which has been observed in other tick species (Kotsyfakis et al., 2015; Lu et al., 2026). Collectively, these results support a conserved midgut transcriptome triggered by blood feeding in hard ticks.

**Figure 6.**
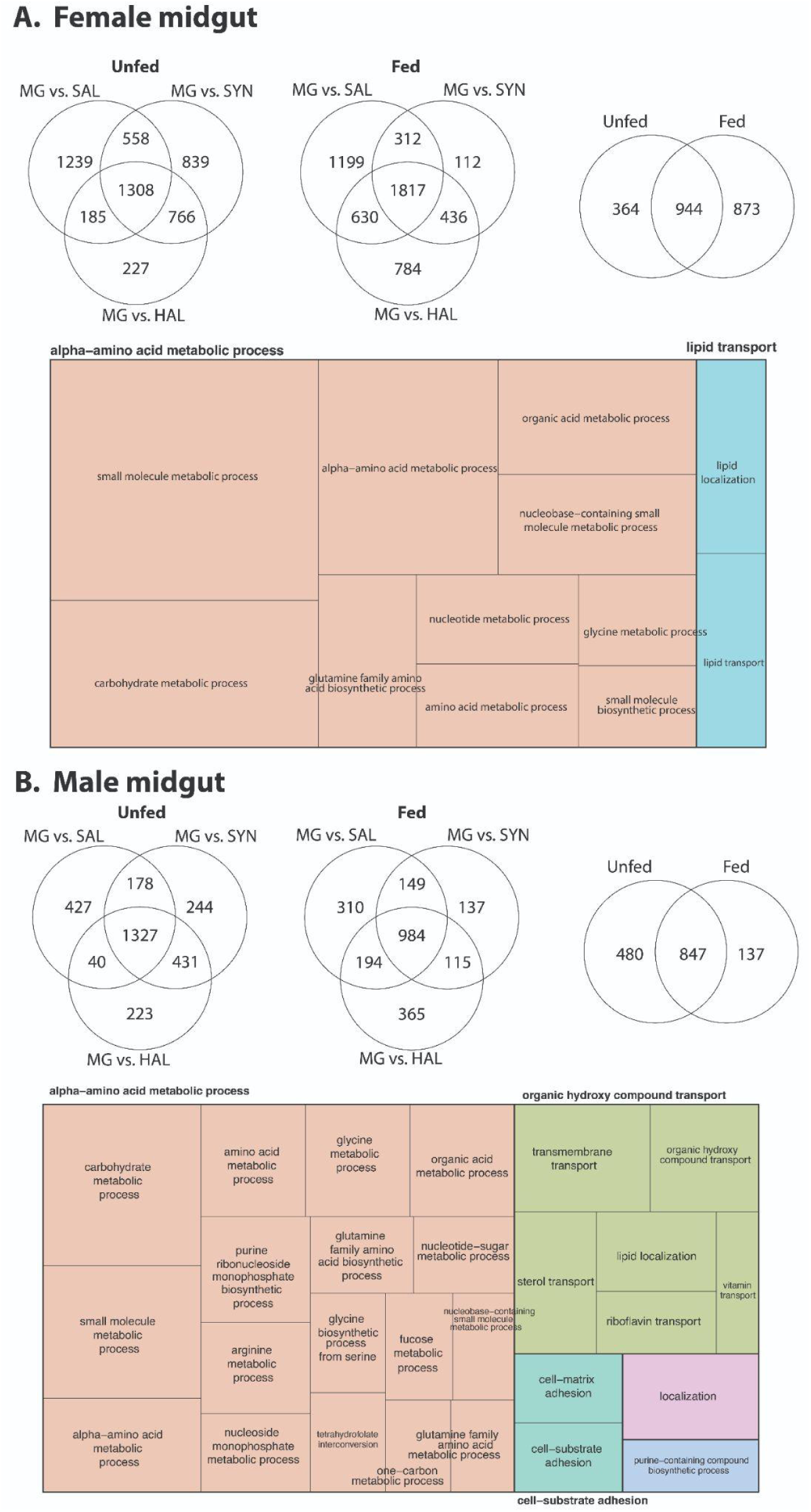
Transcriptomic responses of midgut tissues in *D. andersoni*. RNA-seq–based differential expression analyses of female (A) and male (B) midguts from unfed and fed *D. andersoni*. Venn diagrams depict the number of differentially expressed genes (DEGs) in midgut (MG) tissue relative to salivary glands (SAL), synganglion (SYN), and hemolymph (HAL), highlighting shared and tissue-specific transcriptional responses before and after blood feeding. Overlapping regions indicate DEGs common across comparisons, whereas non-overlapping regions represent genes that are uniquely regulated. Treemap visualizations summarize enriched Gene Ontology (GO) biological processes among midgut-enriched DEGs, with rectangle size proportional to the number of genes assigned to each category. Complete results for female and male ticks are in Table S6 and S7, respectively.

Comparisons of salivary glands with midgut, synganglion, and Haller’s organ also reveal extensive tissue-specific gene expression (Fig. 7, Table S8-9), emphasizing the specialized role of salivary glands in mediating host–tick interactions (Kazimírová and Štibrániová, 2013; Šimo et al., 2017). The pronounced enrichment of enzyme regulator activity, serine-type endopeptidase inhibitor activity, and cytokine/chemokine-binding functions is consistent with the canonical role of tick saliva in counteracting host hemostatic, inflammatory, and immune defenses (Esteves et al., 2017; Kazimírová and Štibrániová, 2013; Medina et al., 2022; Perner et al., 2018; Šimo et al., 2017). These functional categories are hallmarks of tick salivary secretions and support the idea that successful prolonged feeding relies on the coordinated deployment of protease inhibitors and immunomodulatory proteins during feeding. Blood feeding further amplifies these patterns, indicating that salivary gland gene expression is coupled to feeding state and likely reflects the need to modify the salivary cocktail as feeding progresses continually (de Castro et al., 2016; Martins et al., 2021; Medina et al., 2022). Notably, the enrichment of ribosomal structural components and translation-related activities in blood-fed salivary glands suggests a substantial increase in protein synthesis capacity, consistent with sustained secretion during the multi-day blood meal. Tick salivary glands function as highly responsive secretory organs with transcriptional plasticity during blood feeding.

**Figure 7.**
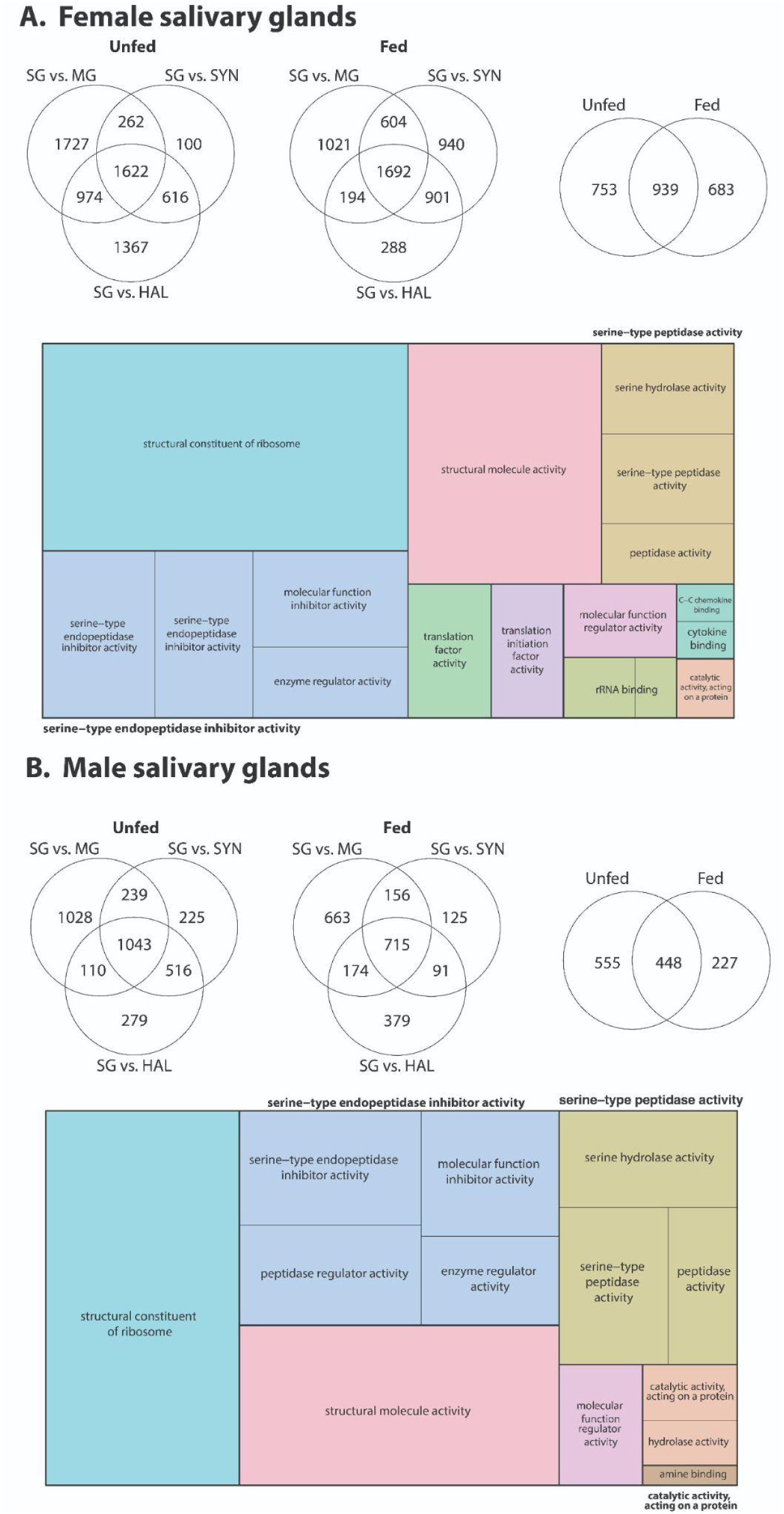
Transcriptomic responses of salivary glands in *D. andersoni* before and after blood feeding. RNA-seq differential expression analyses were conducted on salivary glands (SG) from unfed and fed female (A) and male (B) *D. andersoni.* Venn diagrams show the number of differentially expressed genes (DEGs) in salivary glands relative to midgut (MG), synganglion (SYN), and hemolymph (HAL), highlighting both SG-specific and shared transcriptional profiles before and after blood feeding. Overlapping regions indicate DEGs common among tissue comparisons, whereas non-overlapping regions represent uniquely enriched salivary gland transcripts. Treemap visualizations summarize enriched Gene Ontology (GO) molecular function categories among salivary gland–enriched DEGs. Complete results for female and male ticks are in Table S8 and S9, respectively.

Our results indicate that blood feeding induces pronounced transcriptional reprogramming in the synganglion of *D. andersoni*, with strong enrichment of genes involved in neuronal signaling, ion transport, and regulatory processes, and with broadly similar but not identical patterns between females and males (Fig. 8, Table S10-11). In both sexes, comparisons of synganglion tissue before and after feeding reveal increased G protein–coupled receptor signaling, cyclic nucleotide biosynthetic and metabolic processes, and monoatomic (particularly potassium) ion transmembrane transport, consistent with heightened neurophysiological activity required to integrate sensory inputs and coordinate feeding-associated behaviors that have been observed in other ticks (Bissinger et al., 2011; Lees et al., 2010; Rispe et al., 2022; Zhu et al., 2016). Enrichment of terms related to cellular responses to stimuli, signal transduction, and biological regulation further suggests that the synganglion functions as a dynamic regulatory hub that responds to the physiological demands imposed by blood ingestion. In addition, feeding-associated changes in cell projection organization, cell morphogenesis, neuron differentiation, and cell adhesion imply ongoing neural plasticity or remodeling that may support altered sensory processing or motor control during and after feeding. Overall, these data highlight the central role of the tick synganglion in integrating metabolic state with neural signaling pathways and underscore how blood feeding reshapes the molecular landscape of the tick nervous system.

**Figure 8.**
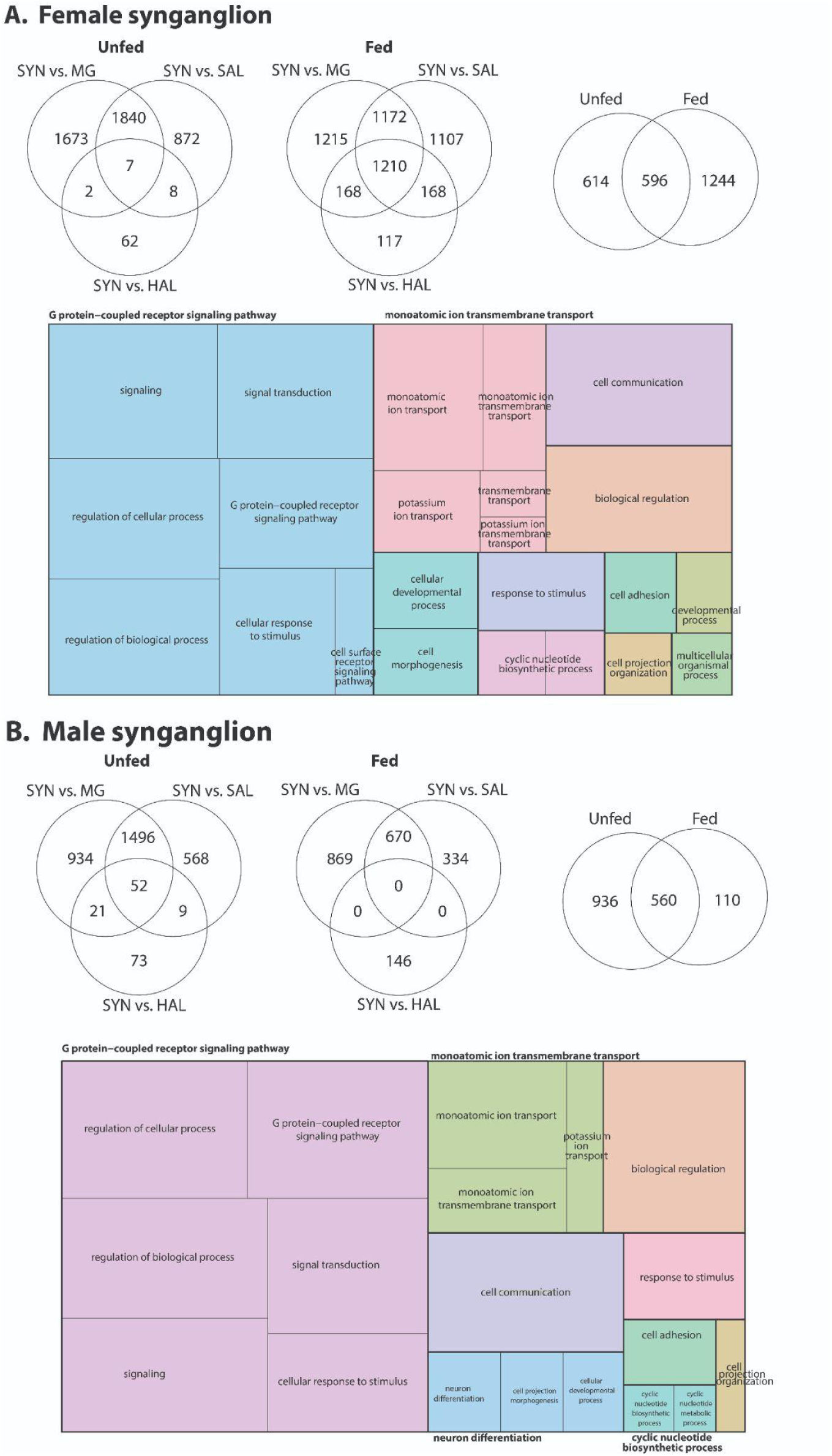
Transcriptomic responses of the synganglion in *D. andersoni* before and after blood feeding. RNA-seq differential expression analyses were performed on the synganglion (SYN) from unfed and fed female (A) and male (B) *D. andersoni*. Venn diagrams illustrate the number of differentially expressed genes (DEGs) in the synganglion relative to midgut (MG), salivary glands (SAL), and hemolymph (HAL), highlighting SYN-enriched and shared transcriptional profiles before and after blood feeding. Additional Venn diagrams compare DEGs between unfed and fed states, indicating both conserved and feeding-responsive neural transcripts. Treemap visualizations summarize enriched Gene Ontology (GO) biological processes among synganglion-enriched DEGs. Complete results for female and male ticks are in Table S10 and S11, respectively.

### Ovary and testes expression analysis

To characterize the molecular programs underlying reproduction in *D. andersoni*, we analyzed transcriptional data from ovaries and testes (Fig. 9, Table S12). We compared their transcriptional profiles to those of non-reproductive tissues, including salivary glands, synganglion, midgut, and hemolymph. Both reproductive tissues exhibited highly distinct, tissue-specific gene expression signatures, with thousands of DEGs uniquely enriched in ovaries or testes, indicating pronounced sexual and functional specialization of the tick reproductive system.

**Figure 9.**
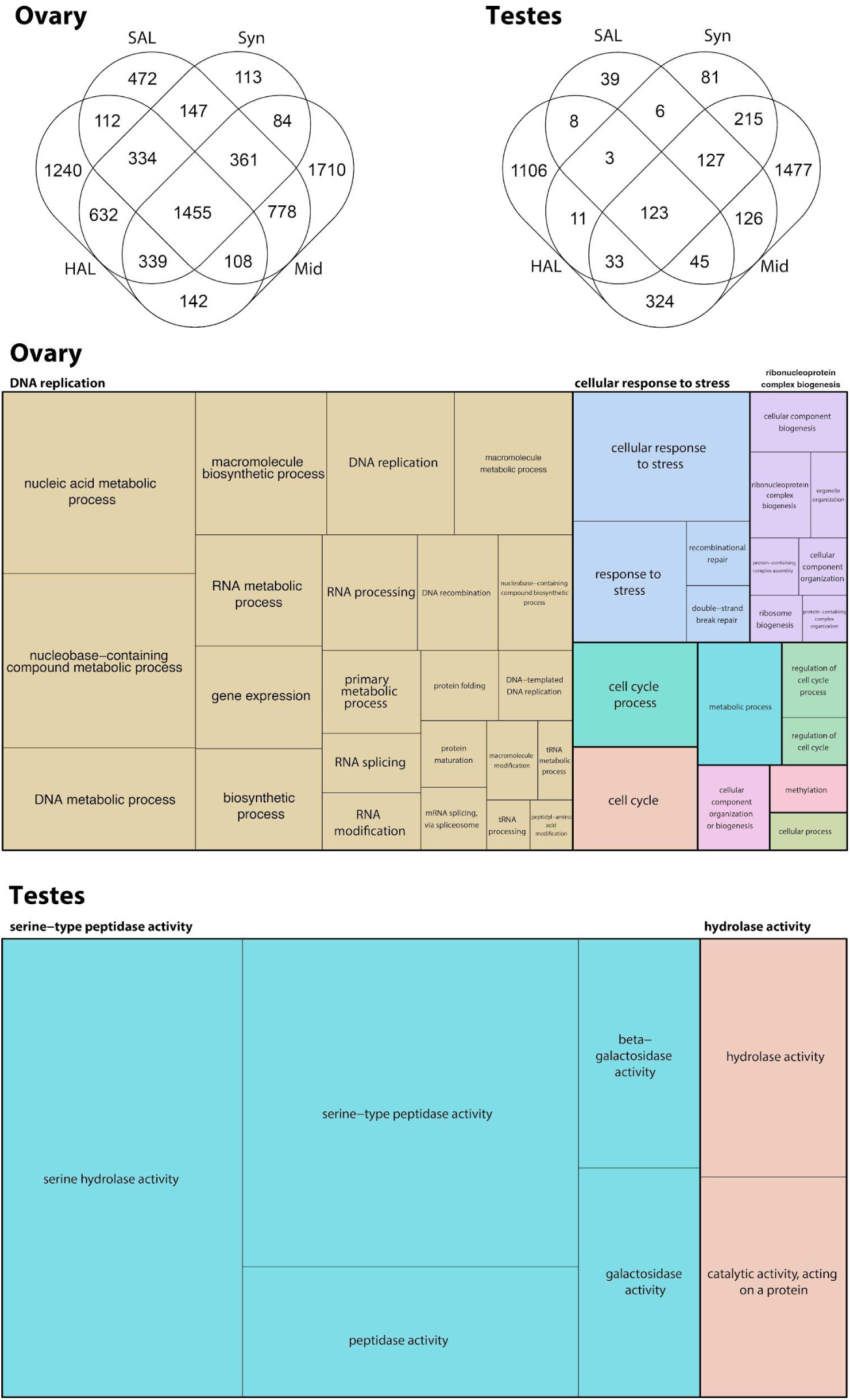
Transcriptomic profiles of reproductive tissues in *D.* andersoni. RNA-seq differential expression analyses were performed on the ovary and testes of *D. andersoni* to characterize sex-specific reproductive transcriptional programs. Venn diagrams illustrate the number of differentially expressed genes (DEGs) enriched in ovaries or testes relative to non-reproductive tissues, including salivary glands (SAL), synganglion (SYN), midgut (MID), and hemolymph (HAL), highlighting both tissue-specific and shared gene expression patterns. Treemap visualizations summarize enriched Gene Ontology (GO) categories among reproductive tissue–enriched DEGs. Complete results for female and male ticks are in Table S12 and S13, respectively.

Ovarian transcriptomes were dominated by genes associated with DNA replication, DNA recombination and repair, RNA processing and splicing, ribosome biogenesis, and macromolecule biosynthetic and metabolic processes. Enrichment of cell cycle regulation, protein folding and maturation, and ribonucleoprotein complex biogenesis further reflects the intense biosynthetic and proliferative demands of oogenesis (Cossío-Bayúgar et al., 2024). The strong representation of DNA-templated transcription, nucleic acid metabolism, and RNA modification pathways suggests sustained transcriptional activity that is required to support germline maintenance, meiotic progression, and the provisioning of maternal transcripts to developing embryos. Similar ovarian expression profiles have been reported in ticks (Cossío-Bayúgar et al., 2024; Wang et al., 2021; Zhao et al., 2021), in which ovary-enriched transcriptomes are characterized by elevated DNA replication, RNA processing, and ribosomal pathways, indicating that these core reproductive programs are highly conserved across hard ticks.

In contrast, testes exhibited a markedly different transcriptional landscape, with DEGs strongly enriched for molecular functions related to proteolysis and enzymatic activity, including serine-type peptidase activity, serine hydrolase activity, and broader peptidase and catalytic activities acting on proteins (Fig 9, Table S13). The prominence of these pathways is consistent with roles in spermatogenesis, sperm maturation, and seminal fluid production (Avila et al., 2010; Meibers et al., 2019), processes that rely heavily on proteolytic remodeling and post-translational regulation. Comparable enrichment of serine proteases and hydrolases has been observed in male transcriptomes of other ticks (Edwards et al., 2025; Gulia-Nuss et al., 2016; Meibers et al., 2019), suggesting that protease-driven regulation of male reproductive function is a conserved feature among ixodid ticks.

### Orthology and transcription factor analyses

To place the *D. andersoni* genome and transcriptome in an evolutionary and regulatory context, we combined orthology inference, transcription factor (TF) identification, and analyses of TF binding motif enrichment with tissue-specific RNA-seq data, with particular emphasis on reproductive tissues (Fig. 10). Orthofinder analyses revealed that *D. andersoni* shares most of its gene families with other hard ticks, including *I. scapularis*, while retaining lineage-specific expansions (Table S14). These shared orthogroups encompass core cellular and metabolic functions, whereas tick-specific and *Dermacentor*-enriched families likely reflect adaptations associated with hematophagy, long life cycles, and reproduction. Importantly, a significant proportion of ortholog expansion was associated with transcription factors (Fig. 10). The orthology structure mirrors patterns observed in other ixodid genomes (Jia et al., 2020), supporting the view that tick genomes are highly conserved in gene content but diversified through gene family expansion.

**Figure 10.**
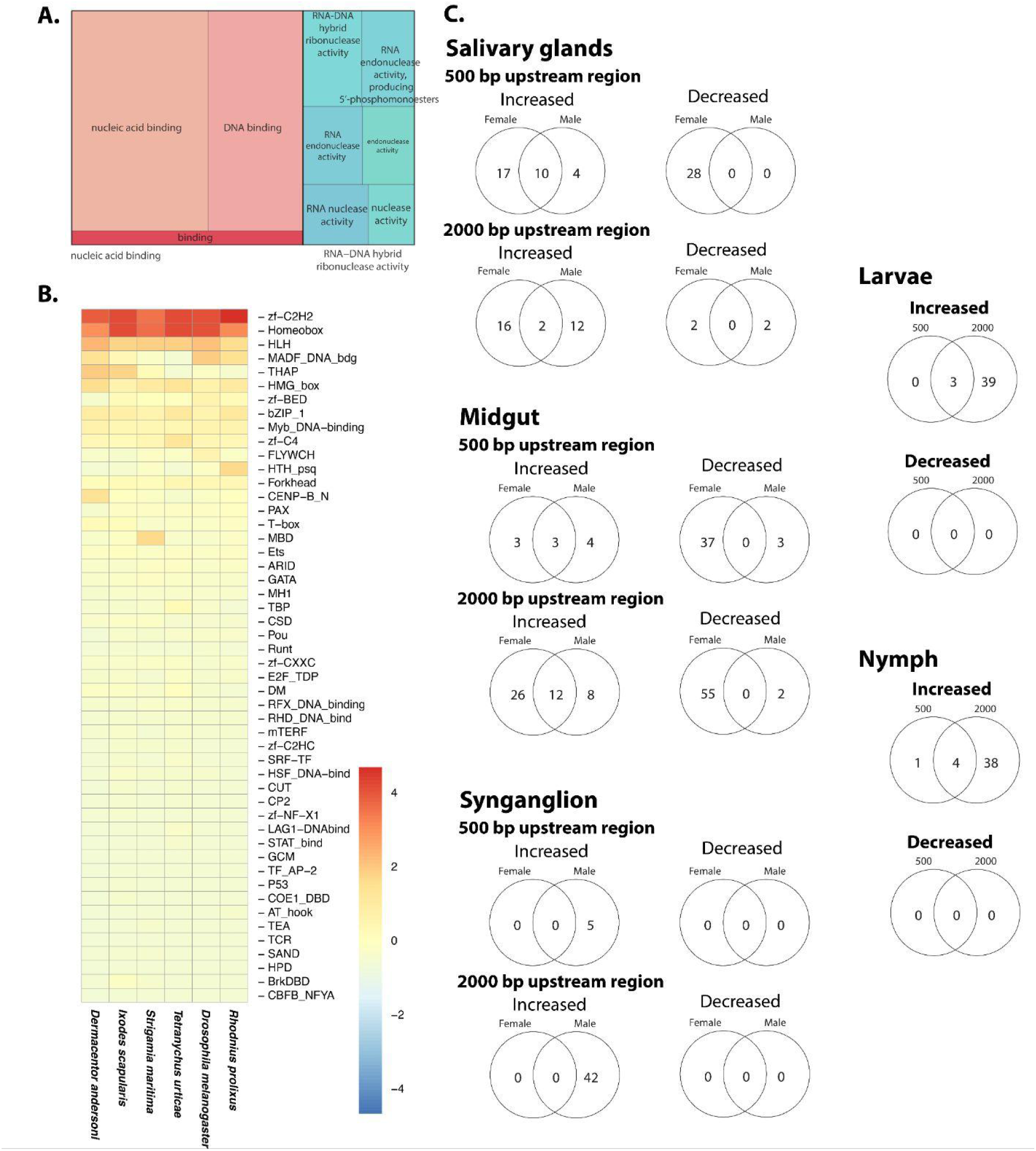
Comparative orthology, transcription factor repertoire, and transcription factor binding site enrichment in *D. andersoni*. (A) Functional annotation of predicted *D. andersoni* expanded orthology groups based on Orthofinder. (B) Heatmap showing the distribution and relative abundance of TF families in *D. andersoni* compared with other arthropods. Family names at the right indicate Pfam IDs. (C) Transcription factor binding motif enrichment analysis using HOMER in promoter regions (500 bp and 2000 bp upstream of transcription start sites) for genes with increased or decreased expression across tissues and developmental stages in relation to blood feeding. Venn diagrams summarize shared and sex-specific enriched motifs in salivary glands, midgut, and synganglion, as well as stage-specific enrichment in larvae and nymphs across different feeding states. Complete results for female and male ticks are in Table S14 and S23, respectively.

Genome-wide identification of genes encoding transcription factors in *D. andersoni* revealed a diverse and canonical TF repertoire, dominated by zinc-finger C2H2, homeobox, helix–loop–helix (HLH), MADF, bZIP, forkhead, GATA, MYB, ETS, and nuclear receptor–associated DNA-binding domains (Fig. 10, Table S15). Comparative analyses across arthropods showed that the relative abundance of these TF families in *D. andersoni* closely resembles that of other ticks. This pattern is consistent with previous reports that ticks retain expanded zinc-finger and homeobox TF complements, likely supporting complex developmental programs, prolonged life stages, and extensive tissue remodeling during feeding and reproduction (Panfilio et al., 2019; Thomas et al., 2020).

Assessment of putative TF binding in promoter regions further highlighted regulatory divergence among tissues and developmental stages (Fig. 10, Tables S16-22). TF binding enrichment patterns differed depending on promoter window size (500 bp versus 2000 bp upstream), tissue type, sex, and feeding state, indicating that regulatory control in ticks operates across multiple spatial scales and is dynamically modulated. Notably, reproductive tissues exhibited stronger and more consistent TF-binding enrichment than somatic tissues, supporting the hypothesis that reproduction is under tight transcriptional control. In larvae and nymphs, TF binding enrichment patterns shifted with development, consistent with stage-specific regulatory programs that precede and accompany blood feeding, as reported in other tick species.

Together, these integrated analyses demonstrate that *D. andersoni* shares a highly conserved core gene and TF repertoire with other ixodid ticks with expansions in specific families while exhibiting pronounced tissue- and sex-specific regulatory specialization. The strong correspondence between *D. andersoni* and other tick species suggests that much of tick biology is governed by conserved regulatory frameworks. In contrast, species-specific adaptations are likely mediated through differential deployment of shared TFs and gene families. By linking orthology, transcription factor diversity, and reproductive transcriptomics, this study provides a comprehensive view of how genome content and regulatory architecture interact to shape tick development and reproduction, offering a foundation for comparative studies and for targeting regulatory pathways relevant to tick control and vector competence.

### Identification of the sex chromosome

*Dermacentor andersoni* follows an XO-sex determination system, where females have two X chromosomes, while males only have one (Oliver, 1972). To evaluate the sex chromosome among 11 chromosomes in *D. andersoni*, we sequenced whole genomes of seven adult males and seven adult females and inspected relative coverage across all chromosomes. We found that relative coverage of the longest chromosome (CM094977.1) in females was almost twice the coverage of males (Figure 11). This coverage difference was consistent between nymphs and adults, indicating that difference in coverage was a result of sex, as there was no significant difference in relative coverage at this chromosome between nymphs and adults (F1,26=1.69, p=0.2, Table S24), but there was between males and females (F1,26=3745, P<0.001, Table S24). This corroborates with karyotyping studies that the sex chromosome is the largest chromosome and is slightly longer than the largest autosome (Oliver 1972). Our results conclusively identify that the first chromosome, which is also the longest, is the sex chromosome in *D. andersoni*.

**Figure 11.**
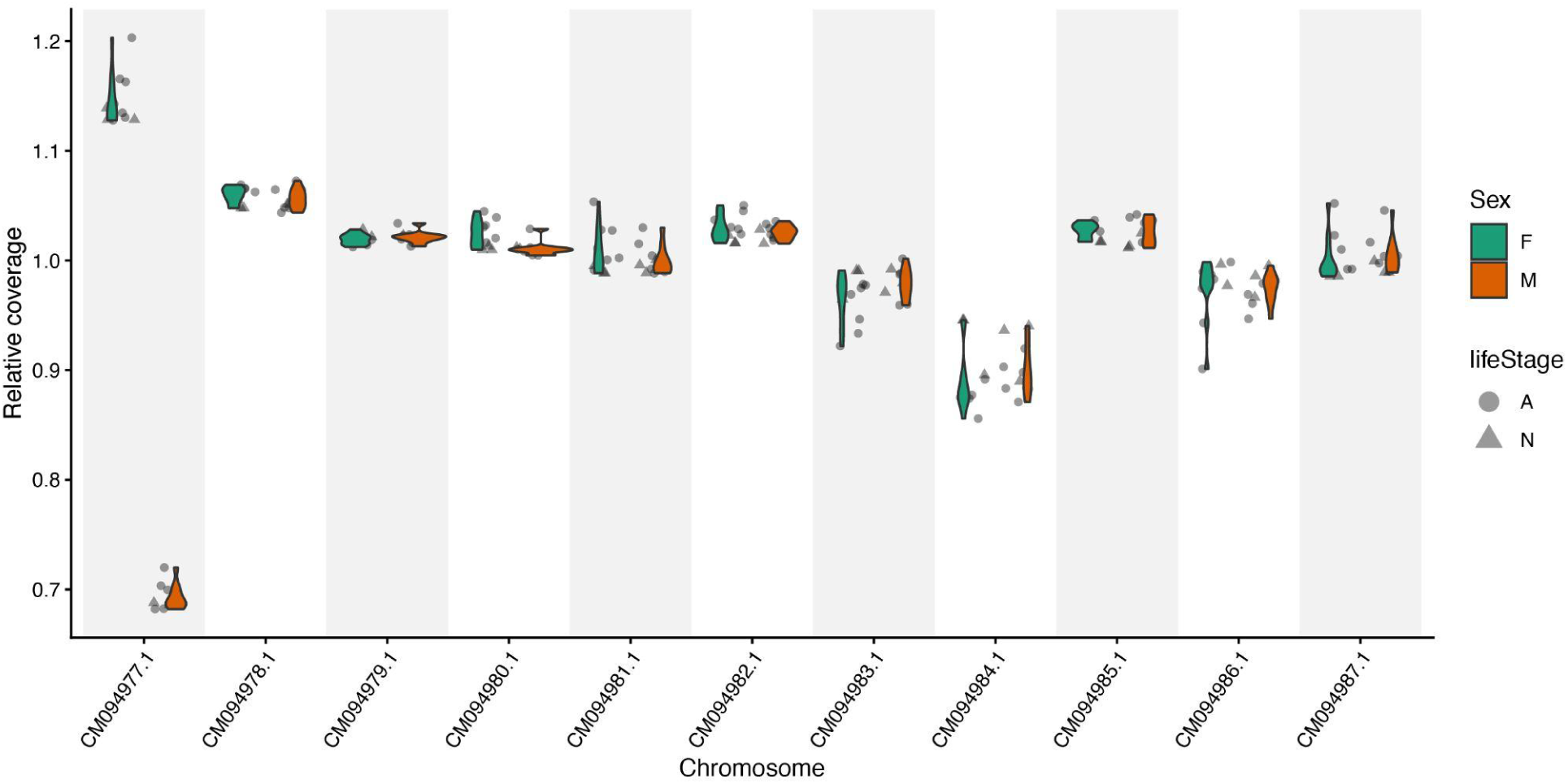
Relative sequencing coverage across chromosomes. The relative coverage across *D. andersoni* chromosomes from sequence data generated from 14 adults and 6 nymphs reveals substantially lower coverage on the first and largest chromosome, CM094977. There was no difference in coverage between life stages, but there was between sexes (see Results and Discussion).

### Summary

Here, we present a high-quality, chromosome-level genome assembly for *D. andersoni*, the Rocky Mountain wood tick, together with an extensive and deeply sampled RNA-seq dataset encompassing multiple developmental stages, sexes, tissues, and feeding states. The genome assembly exhibits high contiguity and completeness, as evidenced by the recovery of the vast majority of benchmarking universal single-copy orthologs, providing strong confidence in its accuracy and suitability for downstream analyses. This assembly is among the most comprehensive genomic resources currently available for hard ticks (De et al., 2023; Jia et al., 2020). It enables robust investigations of genome organization, gene family evolution, and the genetic basis of key life-history traits.

The transcriptomic datasets capture dynamic patterns of gene expression across eggs, larvae, nymphs, and adults, as well as across tissues including midgut, salivary glands, nervous tissue, and reproductive organs, with particular emphasis on transcriptional changes associated with blood feeding. These data reveal pronounced stage-, tissue-, sex-, and feeding-dependent shifts in metabolic, regulatory, and signaling pathways, highlighting the extensive transcriptional reprogramming that accompanies hematophagy.

Together, these genomic and transcriptomic resources provide unprecedented resolution into the molecular mechanisms governing wood tick development, physiology, and host–pathogen interactions. Importantly, this integrated genomic and transcriptomic framework enables the identification of candidate genes and pathways involved in blood meal acquisition, digestion, immune modulation, and pathogen transmission, all of which are central to vector competence. Beyond advancing fundamental understanding of tick biology, these datasets provide a critical platform for comparative genomics across arthropod vectors and for the development of novel tick- and disease-control strategies targeting conserved or tick-specific molecular processes.

## Acknowledgements

This study was partially supported by shared computational resources by the National Institute of Allergy and Infectious Diseases of the National Institutes of Health under Award Numbers R01AI148551 (JBB) and R21AI166633 (JBB), and USDA-ARS CRIS# 2090-32000-043 (PO). MTW and XC were supported by NIH grants U24HG013078 and P30AR070549. HY was supported by the Eyes High Postdoctoral Fellowship at the University of Calgary and JK was supported by the Margaret Gunn Animal Endowment Fund. This work was supported by the United States Department of Agriculture, Agricultural Research Service (USDA-ARS) with appropriated funding to the USDA-ARS Ag100Pest Initiative, USDA-ARS Tropical Pest Genetics and Molecular Biology Research Unit Project 2040-30400-003-000D, and the USDA-ARS SCINet Project 0500-00093-001-00-D. We also acknowledge the support of the Natural Sciences and Engineering Research Council of Canada (NSERC), funding reference number RMS21-73779779 [Cette recherche a été financée par le Conseil de recherches en sciences naturelles et en génie du Canada (CRSNG), numéro de référence RMS21-73779779]. We also acknowledge the outstanding technical expertise of Shelby M. Jarvis, Kathleen L. Mason, and Gavin Scoles. All opinions expressed in this paper are the author’s and do not necessarily reflect the policies and views of USDA or NIH. Mention of trade names or commercial products in this publication is solely for the purpose of providing specific information and does not imply recommendation or endorsement by USDA or NIH. USDA and NIH are equal opportunity providers and employers.

## Data availability

All datasets are available through the NCBI (BioProject PRJNA936406).

## Supplementary figures

**Figure S1 -** Weighted Gene Co-expression Network Analysis (WGCNA) was used to identify modules of co-expressed genes across multiple tissues and developmental stages of *Dermacentor andersoni*.

## Supplementary Materials

**Materials S1 -** List of transcripts associated with specific modules based on Figure S1.

## Supplementary tables

**Table S1 -** Summary of predicted tRNA genes from *Dermacentor andersoni* in comparison to other arthropods

**Table S2 -** Genes with enriched transcript levels in *Dermacentor andersoni* eggs compared to other development stages. Significance determined by stage-specific comparisons with DESeq2 (FDR, P < 0.05).

**Table S3 -** Genes with enriched transcript levels in *Dermacentor andersoni* adults compared to other development stages. Significance determined by stage-specific comparisons with DESeq2 (FDR, P < 0.05).

**Table S4 -** Genes with enriched transcript levels in unfed *Dermacentor andersoni* nymphs and larvae compared to blood fed individuals. Significance determined by stage-specific comparisons with DESeq2 (FDR, P < 0.05).

**Table S5 -** Genes with enriched transcript levels in fed *Dermacentor andersoni* nymphs and larvae compared to unfed individuals. Significance determined by stage-specific comparisons with DESeq2 (FDR, P < 0.05).

**Table S6 -** Genes with enriched transcript levels in the midgut of *Dermacentor andersoni* females compared to other tissues. Significance determined by stage-specific comparisons with DESeq2 (FDR, P < 0.05).

**Table S7 -** Genes with enriched transcript levels in the midgut of *Dermacentor andersoni* males compared to other tissues. Significance determined by stage-specific comparisons with DESeq2 (FDR, P < 0.05).

**Table S8 -** Genes with enriched transcript levels in salivary glands of *Dermacentor andersoni* females compared to other tissues. Significance determined by stage-specific comparisons with DESeq2 (FDR, P < 0.05).

**Table S9 -** Genes with enriched transcript levels in salivary glands of *Dermacentor andersoni* males compared to other tissues. Significance determined by stage-specific comparisons with DESeq2 (FDR, P < 0.05).

**Table S10 -** Genes with enriched transcript levels in the synganglion of *Dermacentor andersoni* females compared to other tissues. Significance determined by stage-specific comparisons with DESeq2 (FDR, P < 0.05).

**Table S11 -** Genes with enriched transcript levels in the synganglion of *Dermacentor andersoni* males compared to other tissues. Significance determined by stage-specific comparisons with DESeq2 (FDR, P < 0.05).

**Table S12 -** Genes with enriched transcript levels in ovaries of *Dermacentor andersoni* females compared to other tissues. Significance determined by stage-specific comparisons with DESeq2 (FDR, P < 0.05).

**Table S13 -** Genes with enriched transcript levels in testes of *Dermacentor andersoni* males compared to other tissues. Significance determined by stage-specific comparisons with DESeq2 (FDR, P < 0.05).

**Table S14 -** Summary of orthogroup statistics for *D. andersoni*. The table reports per-species metrics, including the total number of genes assigned to orthogroups and the percentage of genes contained within orthogroups.

**Table S15 -** Identification and characterization of putative transcription factors (TFs). Amino acid sequences of predicted protein-coding genes were scanned for conserved DNA-binding domains (DBDs) to identify putative TFs.

**Table S16 -** HOMER-predicted transcription factor binding sites based on RNA-seq data in adult female midguts before and after blood feeding. Analyses were performed using promoter regions defined as either 500 bp or 2,000 bp upstream of annotated transcription start sites, with enriched motifs reported along with their associated transcription factors and statistical significance values.

**Table S17 -** HOMER-predicted transcription factor binding sites based on RNA-seq data in adult female salivary glands before and after blood feeding. Analyses were performed using promoter regions defined as either 500 bp or 2,000 bp upstream of annotated transcription start sites, with enriched motifs reported along with their associated transcription factors and statistical significance values.

**Table S18 -** HOMER-predicted transcription factor binding sites based on RNA-seq data in adult female synganglion before and after blood feeding. Analyses were performed using promoter regions defined as either 500 bp or 2,000 bp upstream of annotated transcription start sites, with enriched motifs reported along with their associated transcription factors and statistical significance values.

**Table S19 -** HOMER-predicted transcription factor binding sites based on RNA-seq data in adult male midguts before and after blood feeding. Analyses were performed using promoter regions defined as either 500 bp or 2,000 bp upstream of annotated transcription start sites, with enriched motifs reported along with their associated transcription factors and statistical significance values.

**Table S20 -** HOMER-predicted transcription factor binding sites based on RNA-seq data in adult male salivary glands before and after blood feeding. Analyses were performed using promoter regions defined as either 500 bp or 2,000 bp upstream of annotated transcription start sites, with enriched motifs reported along with their associated transcription factors and statistical significance values.

**Table S21 -** HOMER-predicted transcription factor binding sites based on RNA-seq data in adult male synganglion before and after blood feeding. Analyses were performed using promoter regions defined as either 500 bp or 2,000 bp upstream of annotated transcription start sites, with enriched motifs reported along with their associated transcription factors and statistical significance values.

**Table S22 -** HOMER-predicted transcription factor binding sites based on RNA-seq data from larvae before and after blood feeding. Analyses were performed using promoter regions defined as either 500 bp or 2,000 bp upstream of annotated transcription start sites, with enriched motifs reported along with their associated transcription factors and statistical significance values.

**Table S23 -** HOMER-predicted transcription factor binding sites based on RNA-seq data in nymphs before and after blood feeding. Analyses were performed using promoter regions defined as either 500 bp or 2,000 bp upstream of annotated transcription start sites, with enriched motifs reported along with their associated transcription factors and statistical significance values.

**Table S24** - Results comparing relative coverage on chromosome one between life stages and sexes.

