## Supplementary figures and images for "Genome sequencing and multi-stage, blood-feeding, and tissue-specific transcriptome atlas of the Rocky Mountain wood tick provide a critical resource for this vector"

### Supplemental Figure 1

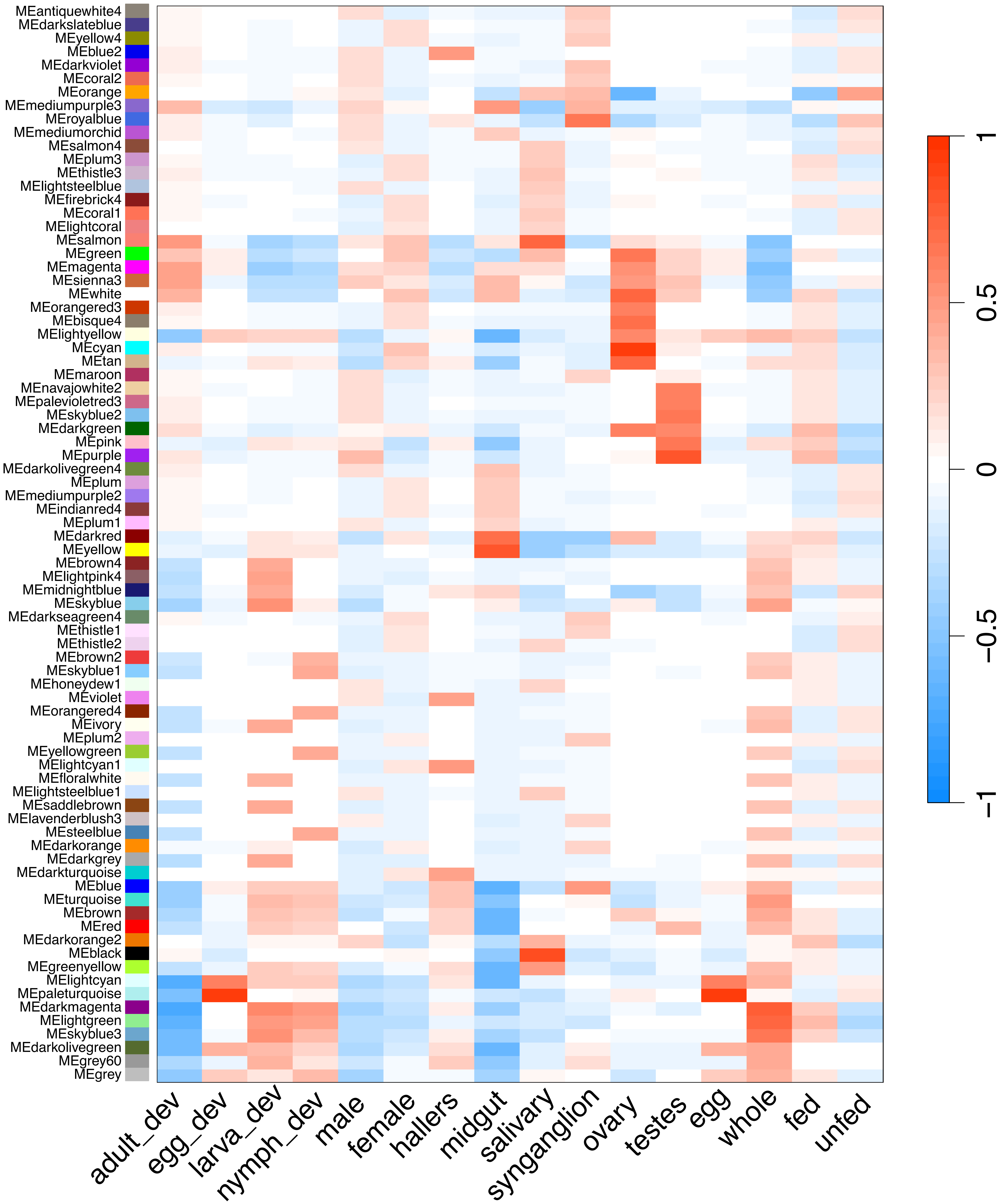
